# Marine medaka responds differently to dioxin compared with its close freshwater relative, the Japanese medaka: the AHR molecular mechanism

**DOI:** 10.1101/2024.12.01.626272

**Authors:** Wanglong Zhang, Yanjiao Zong, Ruize Sun, Zhenhong Xue, Wenhui Wan, Anran Ren, Yongchao Ma, Wenjing Tian, Renjun Wang

**Author notes:** Address correspondence to Dr. Wanglong Zhang, College of Life Sciences, Qufu Normal University, Qufu, Shandong 273165, China. Renjun Wang, College of Life Sciences, Qufu Normal University, Qufu, Shandong 273165, China. Wanglong Zhang and Yanjiao Zong contributed equally as co-first authors.

## Abstract

The global water pollution now calls for precise risk assessment of chemicals, e.g., dioxins and the emerging dioxin-like compounds (DLCs). The freshwater and marine medaka have been widely implemented in the toxicity testing, and perhaps give mechanistic information for comparative biology. The question that ‘will they report equal results due to their close phylogenetic relation’ has been raised, therefore, we explored their physiological and molecular responses to dioxin. As the mediator of the dioxin toxicity, the aryl hydrocarbon receptor (AHR) of marine medaka (*Oryzias melastigma*) has not been functionally characterized and might be species-specific. In terms of sensitivity to dioxin—2,3,7,8-tetrachlorodibenzo-*p*-dioxin (TCDD), the EC_50_ values of omeAHR1a (0.16±0.12 nM), omeAHR1b (2.96±2.96 nM), omeAHR2a (0.44±0.30 nM), and omeAHR2b (9.00±6.88 nM) exhibit marked variations. The omeAHR2a and omeAHR1a display heightened sensitivity compared to the freshwater Japanese medaka (*Oryzias latipes*) counterparts olaAHR2a and olaAHR1a, respectively. The results indicate the in vitro sensitivity of AHR among species can vary by one or two orders of magnitude. Physiologically, the marine medaka (LC_50_: 1.64 ng/L (95% CI: 1.05-2.55 ng/L)) also exhibits a pronounced sensitivity to TCDD than Japanese medaka (LC_50_: 3.42 ng/L (95% CI: 1.37-6.48 ng/L), aligning with the in vitro AHR sensitivity. Further mechanistic investigations using additional ligands and computational modeling reveal that: 1) most of omeAHR2a, olaAHR2a, dreAHR2, and hsaAHR interact with ligands in the affinity order of TCDD > PCB126 > BNF > indole, mirroring their AHR transactivation potency, but the docking poses and dynamics can vary; 2) one AHR subform’s high sensitivity to dioxin—TCDD may extend to DLCs but not to other types of ligands. These insights underscore the difference of AHR biology among species even the close relative species, and finger out the necessity for meticulous consideration when evaluating the toxicity of compounds and when extending predictive toxicity assessments to more species.

## 1. Introduction

In the era of industrialization and rapid urbanization, the integrity of water resources has become a focal point of global concern. The ubiquitous presence of water pollution, exacerbated by the influx of novel chemical substances, poses a multifaceted threat to ecology ^1–3^ and human health ^4, 5^. Among these contaminants, dioxins and dioxin-like compounds (DLCs) stand out due to their potent toxicity and ability to interact with the aryl hydrocarbon receptor (AHR), a critical regulator of cellular responses to xenobiotics. This interaction triggers a cascade of biological events leading to adverse health effects, underscoring the urgency for rigorous risk assessment strategies. Over recent years, new DLCs have been discovered in the environment, demonstrating significant toxic potential by interacting with the AHR, such as seen with polyhalogenated carbazoles (PHCZs) and polychlorinated diphenyl sulfides (PCDPSs) ^6–8^.

The AHR is a transcription factor within the basic helix-loop-helix/Per-ARNT-Sim (bHLH-PAS) family in diverse taxa, and acts as a ‘sensor’ for dioxins and DLCs ^9–11^. Under the canonical activation model, the AHR, once bound with dioxins in the cytoplasm, is activated and moves to the nucleus to bind the dioxin-response element (DRE), triggering the expression of target genes with the assistance of its partner—aryl hydrocarbon receptor translocator (ARNT). The extent of this transactivation is directly related to the expression of target genes and can predict the toxic effects. However, the activation process can vary significantly due to factors such as the ligand’s binding affinity, the selectivity of specific AHR subforms, DRE binding efficiency, and transactivation efficiency, all of which can influence the overall outcome ^12–15^. The World Health Organization (WHO) has reported the toxic equivalent factor (TEF) of numerous dioxins and dioxin-like PCBs for three major groups of species: birds, fish, and mammals (humans) ^16^. This highlights the varying risks that dioxins pose to different species. Understanding these risks can offer further insights into the ecotoxicity caused by dioxins and DLCs.

Despite considerable progress in deciphering the AHR pathway in various model species, there remains a paucity of information regarding the functional nuances and sensitivity variations of this receptor across model species (or wildlife), or even between closely related model species inhabiting different aquatic habitats. For risk evaluation and toxicity testing, the Japanese medaka (*Oryzias latipes*), a small freshwater fish native to East Asia, has long been a cornerstone ^17, 18^. Studies involving Japanese medaka have shed light on the mechanisms of toxicity for various pollutants, and contributed to a deep understanding of how these contaminants impact freshwater ecosystems. However, as we turn our gaze towards the vast and largely unexplored realm of marine environments, the potential of a marine equivalent and also relative species, marine medaka (*Oryzias melastigma*) emerges as an intriguing prospect for advancing our knowledge on marine pollution and its consequences ^19^. Understanding the sensitivity discrepancies between these two medaka species may hinge on subtle variations in the constitution, expression, and function of their AHRs, which in turn could be influenced by their respective evolutionary histories and environmental adaptations. For instance, marine organisms often face unique selective pressures, such as higher salinity and exposure to a distinct suite of pollutants, which could drive the evolution of specialized detoxification mechanisms ^20^.

This gap in knowledge of responses to dioxin and DLCs is pertinent when considering the marine and Japanese medaka. This study, therefore, aims to delve into the marine medaka AHRs’ function and mechanism, particularly including phylogeny, expression profiles, sensitivity, and structural basis. Alongside above aspects, the comparison with Japanese medaka and additional species (e.g., zebrafish and human) was conducted. Finally, uncovering the discrepancies between the marine and freshwater medaka would enhance their application in environmental monitoring, toxicity testing, and benefit the conservation of freshwater and marine ecology.

## 2. Materials and Methods

### 2.1 Marine medaka and Japanese medaka embryo exposure

Embryos of marine medaka and Japanese medaka were procured from Yixiyue Biotechnology (Shandong, China). Exposure experiment was conducted according to the standard method of China (standard numbers: GB/T 29764-2013 and HJ 1069-2019) with minor modifications to adopt to the lab resources. Briefly, embryos at 3 dpf were selected for exposure experiment in 24-well plate. For marine medaka, embryos were maintained in freshly prepared seawater and replaced daily, with water temperature at 25 °C, light-dark ratio of 12 h: 12 h, salinity at 31, and pH ranging from 8.15 to 8.20. For Japanese medaka, embryos were under the same temperature and light conditions in the rearing solution contains NaCl-0.10 g, KCl-0.03 g, CaCl_2_-0.04 g, MgSO_4_-0.16 g and deionized water-100 mL (pH = 7.85, DO = 5.58 mg/L) ^21^. The embryos were exposed to a series of concentrations of 2,3,7,8-tetrachlorodibenzo-*p*-dioxin (TCDD) and the control (0.05% (v/v) dimethyl sulfoxide (DMSO); Sigma, St. Louis, MO, USA). Embryos were examined daily for their developmental stage, visible lesions, and death. After 10 days’ exposure, the LC_50_ and hatching rate were calculated according to the methods described in *section 2.9*. All procedures involving the animals were conducted according to the guidelines set by the Biomedical Ethics Committee of Qufu Normal University (approval number: NO. 2024136). Certification of approval can be provided if requested.

### 2.2 AHR pathway phylogenetic analysis

By conducting a targeted keyword search within the NCBI nucleotide database and employing the Basic Local Alignment Search Tool (BLAST), we successfully identified four AHR and two ARNT homologs in marine medaka. Among them, one AHR was designated with a specific nomenclature (omeAHR1b: XM_024288956.2), while the others lacked such designation. The sequences obtained from the NCBI were aligned using Clustal Omega software. For the phylogenetic analysis, we employed a maximum-likelihood (ML) approach based on the Jones-Taylor-Thornton substitution model, using MEGA 11. To ensure robustness, the reliability of the branches was evaluated using 1000 bootstrap replicates. A diverse range of nucleotide sequences from various species were involved in the analysis, including *Oryzias latipes*, *Danio rerio*, *Takifugu rubripes*, *Anabas testudineus*, *Cyprinus carpio*, *Caenorhabditis elegans*, *Rattus norvegicus*, *Homo sapiens*, and *Mus musculus*. The specific NCBI accession numbers for these sequences are detailed in Supplementary **Table S1**. It is important to note that the prefixes ‘ola’, ‘dre’, and similar are derived from the initial letters of the Latin names for the respective species.

### 2.3 AHR pathway expression quantification

The AHR pathway is composed of key genes such as AHR, ARNT, and the aryl hydrocarbon receptor repressor (AHRR). To quantify the expression of these genes, we utilized public RNA-seq data from the NCBI Sequence Read Archive (SRA) database. The data sets encompass a range of marine medaka tissues, the accessions are as follows: brain (SRR1281992, SRR1281993 and SRR12415467), gill (SRR10424556, SRR10424557, SRR10424560 and SRR10424561), liver (SRR17333852, SRR17333855, SRR17333856, SRR17333857, SRR17333863, and SRR17333864), muscle (SRR25458686, SRR25458687, and SRR25458688), testis (SRR18494736, SRR18494737, SRR18494738, SRR18494739, and SRR18494740), and embryo (SRR22250803, SRR22250808, and SRR22250809). The RNA-seq data were processed on the Galaxy server, using the Kallisto tool for gene expression quantification. This involved the use of specific gene sequences with the following accessions: omeAHR1a (XM_024286767.2), omeAHR1b (XM_024288956.2), omeAHR2a (XM_024288918.2), omeAHR2b (XM_024286404.1), omeARNT1 (XM_024267527.2), omeARNT2 (XM_024281904.1), and omeAHRR (XM_024280139.2). The gene expression levels were reported in transcripts per kilobase million (TPM), a metric for assessing gene expression from RNA-seq data.

### 2.4 Expression vector construction of omeAHR and omeARNT

Marine medaka were anesthetized using a solution of 0.03% tricaine methanesulfonate (MS-222). Under ice bath condition, the fish were swiftly processed into a homogenate. The extraction of total RNA from these homogenates was carried out with the aid of a Total RNA Purification Kit (TianGen, Beijing, China). Following this, a First-Strand cDNA Synthesis Kit (Vazyme, Nanjing, China) was utilized to perform reverse transcription of 2 μg of the total RNA, employing oligo-dT primers for the reaction. Using marine medaka cDNA as a template, the complete coding sequences for four omeAHRs and two omeARNTs were successfully amplified by the Phanta Max high-fidelity PCR enzyme (Vazyme). The primers are listed in **Table S2** and the PCR conditions followed those previously described by Karchner et al. ^22^, which are as follows: an initial denaturation at 94°C for 1 minute, followed by 5 cycles of 94°C for 5 seconds and 72°C for 3 minutes, then 5 cycles of 94°C for 5 seconds and 70°C for 3 minutes, and finally 30 cycles of 94°C for 5 seconds and 68°C for 3 minutes. The PCR-generated products were purified using Gel DNA extraction kit (TianGen), and then cloned into the mammalian expression vector—Dual Promoter driven by a CMV promoter (System Biosciences, Palo Alto, CA, USA) using the Seamless Cloning Kit (Beyotime Biotechnology, Shanghai, China). The ligation mixture was transformed into DH5α competent cells, and positive clones were picked. Plasmid was extracted and sequenced by Sangon Biotech (Shanghai, China). The expression constructs of AHR and ARNT of other species (e.g., Japanese medaka, zebrafish, and human) were constructed or reported previously under the similar protocol ^23^.

### 2.5 Functional screening of omeAHR and omeARNT

COS-7 cells, a widely used cell line originating from African Green Monkey kidney fibroblasts transformed with Simian Virus 40 (SV40), were procured from the National Infrastructure of Cell Line Resource in Beijing, China. These cells are well-suited for the expression of foreign genes and the study of AHR transactivation. They were cultivated at 37°C in a 5% CO_2_ environment using Dulbecco’s Modified Eagle Medium (DMEM; Meilunbio, Dalian, China) supplemented with 10% fetal bovine serum (FBS; Sangon Biotech). For the functional screening assay, the potent AHR agonist—TCDD (Wellington, Ontario, Canada; product code: CUS-DD-48-D) was prepared as a 100 μM stock solution in DMSO. In the assay, COS-7 cells were seeded in a 96-well plate at a density of 40,000 cells per well. Transfection was carried out 12 hours later using Lipo8000 transfection reagent (Beyotime, Shanghai, China) and Opti-MEM (Invitrogen, Burlington, ON, USA). The transfection mixture included AHR (30 ng/well) and ARNT (20 ng/well) expression constructs in various combinations (negative control, single construct, or both) along with a luciferase reporter construct pGL3-dcp (50 ng/well), which is based on the cyp1a promoter of the zebrafish ^24^. To normalize the total transfection amount, herring sperm DNA was added to reach 100 ng/well for the respective conditions. Post-transfection, TCDD was added to the medium at the 100 nM working concentration. Each group, including control and TCDD-treated samples, was performed in triplicate within a single assay. The data were presented as the mean ± SD. After a 24-hour exposure to TCDD, the cells were lysed, and their luminescence was measured using a luciferase reporter gene assay kit (Meilunbio) by Biotek Synergy H1 luminometer. The values were normalized to the protein concentrations measured according to the method of Bradford ^25, 26^.

### 2.6 OmeAHR sensitivity determination

We co-transfected COS-7 cells with the same set of constructs: AHR (30 ng) and ARNT (20 ng) expression vectors, and the pGL3-dcp (50 ng) luciferase reporter vector. Post-transfection, after a 12-hour interval, a gradient of TCDD concentrations, spanning from 0.64 pM to 125 nM, was introduced into the culture medium (0.1% v/v DMSO as the solvent control). Following a 24-hour incubation with TCDD, cell lysates were prepared and analyzed using a luciferase reporter gene assay kit (Meilunbio) by Biotek Synergy H1 luminometer. The values were normalized to the protein concentrations measured according to the method of Bradford ^25, 26^. The data were then normalized and fitted to a logistic equation: Y = Bottom + (Top − Bottom) / (1 + 10^˄^ ^((logEC50 − X) × Hillslope)^). In this equation, X corresponds to the logarithm of the TCDD concentration, and Y represents the relative luminescence activity. The parameters EC_20_, EC_50_, and EC_80_ were derived from the curve fitting for each AHR-ARNT combination, encompassing various marine medaka AHR subforms. For species-specific comparison, we also determined the TCDD-induced dose-response curves of AHR of additional species, including Japanese medaka (olaAHR2a) and zebrafish (dreAHR2). Except for those species, when examining the dose-response curves of hsaAHR, we transfected the AHR reporter vector (pGL3-dcp) to HepG2 cells, and determined the luciferase signal with the same protocol like conducted in COS-7 cells. The relative sensitivity (ReS) of each AHR-ARNT pair was calculated by comparing their EC_50_ values to that of dreAHR2, using the formula: ReS = (EC_50_ of dreAHR2 / EC_50_ of interest). The relative potency (ReP) in activating a given AHR is used for the evaluation of a compound’s ability in activating AHR in comparison with TCDD, which was calculated as follows: ReP_compound_ = EC_50_ of TCDD / EC_50_ of compound. Each AHR-ARNT pair’s dose-response curve was determined in triplicate experiments to ensure reliability and reproducibility.

### 2.7 Determination of the AHR transactivation potency of additional ligands

For more comprehensive understanding of inter-species variations, additional AHR ligands were examined for their ability in transactivating AHR of various species (e.g., Japanese medaka, zebrafish, and human), including 3’,4,4’,5-pentachlorobiphenyl (PCB126; CAS: 57465-28-8, Wellington, Ontario, Canada), beta-naphthoflavone (BNF; CAS: 6051-87-2, aladdin, Shanghai, China), and indole (CAS: 120-72-9, Macklin, Shanghai, China). We tested their activation potency on omeAHR2a, olaAHR2a, dreAHR, and hsaAHR using the same method to TCDD described in *section 2.6*. Before testing the compounds, we performed the MTT assay to evaluate cell viability and ensure the compounds did not largely affect the cell viability and possibly the final luciferase result. In brief, COS-7 and HepG2 cells were seeded at a density of 5000 cells per well and allowed to adhere overnight. Subsequently, various concentrations of the compounds (e.g., TCDD, PCB126, BNF, and indole) were added, and the cells were incubated for 28 hours. After aspirating the supernatant, cells were rinsed with PBS and treated with 100 µL of serum-free DMEM followed by the addition of 10 µL of 0.5% MTT solution. Following a 4-hour incubation, the reaction was terminated by removing the medium and adding 150 µL of DMSO to dissolve the formazan crystals. The absorbance at 490 nm was measured using a Biotek Synergy H1 microplate reader.

### 2.8 Modeling the AHR-ligand interaction

The ligand binding domain (LBD) of AHR in marine medaka, Japanese medaka, zebrafish, and human was constructed using homology modeling. The SWISS-MODEL server was employed to construct the models based on the amino acid sequences of the above AHRs. The human AHR LBD structure (PDB id: 7ZUB.1.D), a cryo-EM structure of the indirubin-bound Hsp90-XAP2- AHR complex, was identified as a suitable template by the server’s algorithm. The resulting models were subjected to a series of quality assessments, including Ramachandran plots, GMQE scores, and QMEAN Z-scores, to ensure structural accuracy. The CASTp server was utilized to identify binding cavities within the protein structure, employing a 1.4 Å probe radius. Initial ligand structure for ligands were sourced from PubChem, and molecular docking was conducted using the DockThor server to predict their binding pose to the AHR LBD. The protein-ligand interactions were further analyzed and visualized using the PLIP server. Subsequently, a molecular dynamics simulation of the ligand-AHR complexes was carried out using GROMACS with the CHARMM36 forcefield over a 100 ns period. The binding free energies of these complexes were estimated using the MM-PBSA method, based on 100 frames extracted from the 95 to 100 ns segment of the simulation.

### 2.9 Data statistics

Nonlinear logistic dose-response curve fitting of the luciferase reporter gene assay was conducted using Prism GraphPad 9 (San Diego, CA, USA). The lethal concentrations for 50% mortality (LC_50_) of embryos along with their respective 95% confidence intervals, were determined utilizing the Environmental Protection Agency’s (EPA) BMDS 3.3 Software. Prior to the significance testing, the data underwent an examination for normal distribution and a test for homogeneity of variance. In cases where the data failed to meet the criteria for normal distribution and homogeneity of variance, a nonparametric approach was taken, employing the Kruskal-Wallis test. On the other hand, when the data satisfied the normal distribution and variance homogeneity tests, parametric statistical methods were applied, specifically the one-way ANOVA and t-test. All statistical analyses were executed using both SPSS 26 and Prism GraphPad 9. The results were considered statistically significant at the levels of \**p* < 0.05 and \*\**p* < 0.01.

## 3. Results and Discussion

### 3.1 Marine medaka AHR and ARNT phylogeny

The prevailing theory is that the vertebrate genome experienced two rounds of whole-genome duplications (WGD) around 600 million years ago at the dawn of vertebrate evolution, with subsequent lineage-specific gene losses that can complicate orthology assignments, including the AHR ^27^. Unlike humans and mice, which possess a single AHR gene, teleosts typically exhibit one or two tandem pairs of AHR1 and AHR2 genes, as well as other forms. For instance, zebrafish exhibit dreAHR1a, dreAHR1b, and dreAHR2 ^28^. Interestingly, despite its name, dreAHR1a is actually orthologous to mammalian AHRs and is located on chromosome 16, rather than resembling the typical AHR1a subform in teleosts. The genes dreAHR1b and dreAHR2 are found in tandem on chromosome 22.

Following BLAST analysis and keyword searches of the NCBI database using the reference marine medaka genome ASM292280v2, we identified four AHRs and two ARNTs. As depicted in **Fig. 1A**, marine medaka AHRs can be categorized into the AHR1 and AHR2 subfamilies, with two pairs of AHR1-AHR2 genes located on chromosomes LG21 and LG2, respectively. No additional AHR gene was detected in the marine medaka genome. Marine medaka ARNTs, essential partners for AHR transactivation, were also analyzed phylogenetically. They were found belonging to the ARNT1 and ARNT2 subfamilies (e.g., omeARNT1 and omeARNT2), similar to those in humans, mice, and zebrafish, with two forms present (**Fig. 1B**). Given the diversity of marine medaka AHRs and ARNTs, further bioinformatics and functional studies are warranted to elucidate their roles. Amino acid sequences of the four marine medaka AHRs were aligned with the mouse mmuAHR using Clustal Omega. Similar to the mmuAHR, marine medaka AHRs also conserve functional domains in the N-terminal region, including the bHLH, nuclear localization signal (NLS), nuclear export signals (NES), and PAS domains (**Fig. 2**). Sequence identity analysis revealed varying degrees of similarity between marine medaka and mouse AHR, e.g., omeAHR1a vs. mmuAHR (40.16%), omeAHR1b vs. mmuAHR (39.46%), omeAHR2a vs. mmuAHR (31.01%), and omeAHR2b vs. mmuAHR (31.57%). Similarly, the Japanese medaka olaAHR1a, olaAHR1b, olaAHR2a, and olaAHR2b respectively shows 38.36%, 36.16%, 28.34%, and 27.05% identity. Likewise, zebrafish AHRs exhibit sequence identities of 33.41% for dreAHR1a, 37.88% for dreAHR1b, and 27.85% for dreAHR2 when compare to the mouse mmuAHR. Notably, the sequence identities between marine medaka and Japanese medaka are much greater, e.g., AHR1a (76.82%), AHR1b (84.81%), AHR2a (79.66%), and AHR2b (82.64%). Regarding the distinct region of AHR, the N-terminus (1-468 AA of mmuAHR and the omeAHRs counterparts) of omeAHRs is much conserved and have an average 57.90% identity in comparison with mmuAHR, in contrast the C-terminus of AHRs (469–805 AA) was just 11.41%.

**Figure 1.**
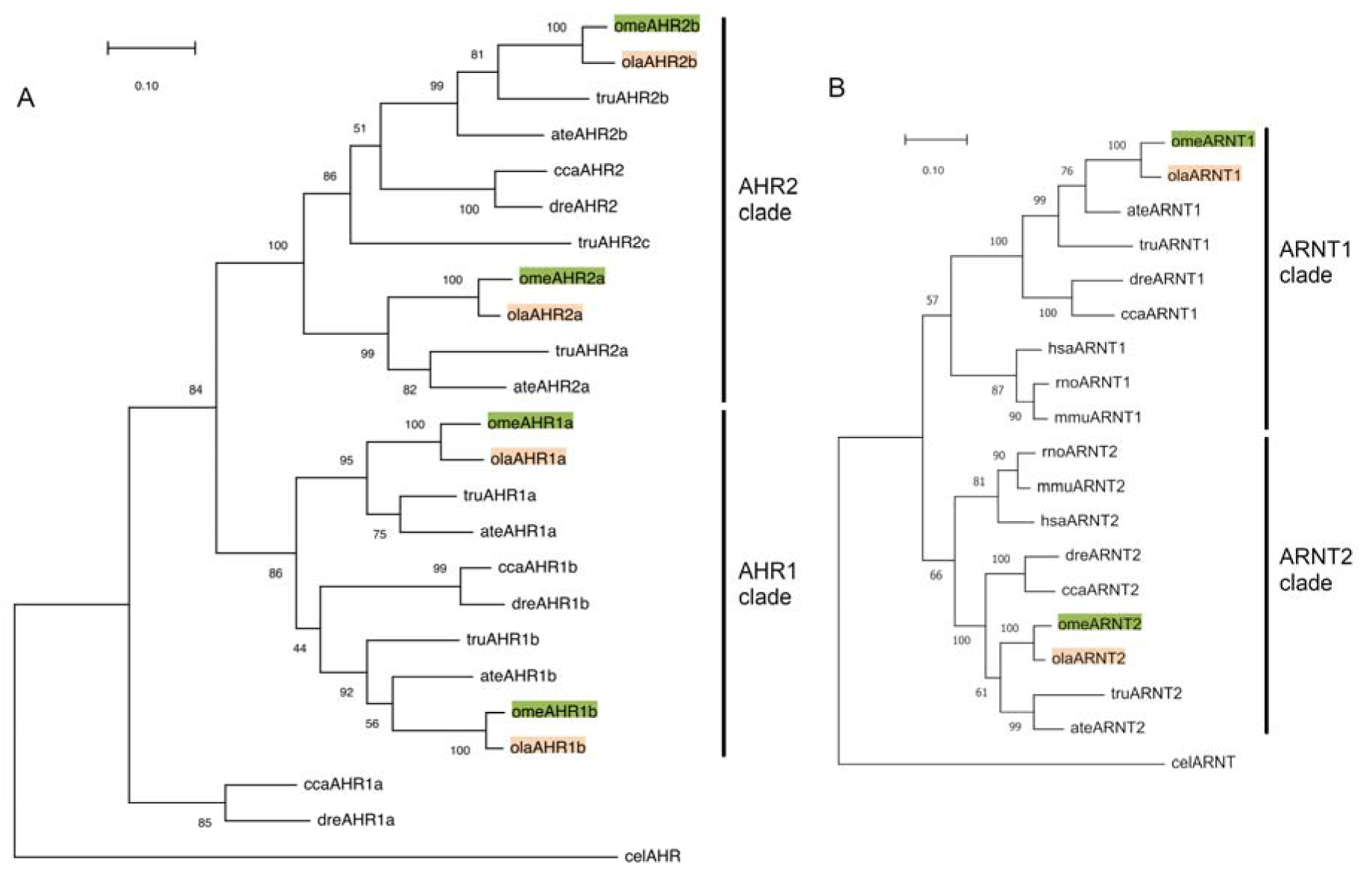
Maximum-likelihood phylogenetic analysis is depicted with node numbers representing bootstrap values derived from 1000 replicates. A) The phylogenetic examination of the AHR gene in marine medaka, and B) The phylogenetic examination of the ARNT gene in marine medaka. The abbreviations correspond to various species: *Oryzias melastigma* (ome), *Oryzias latipes* (ola), *Danio rerio* (dre), *Takifugu rubripes* (tru), *Anabas testudineus* (ate), *Caenorhabditis elegans* (cel), *Homo sapiens* (hsa), *Rattus norvegicus* (rno), *Cyprinus carpio* (cca), and *Mus musculus* (mmu).

**Figure 2.**
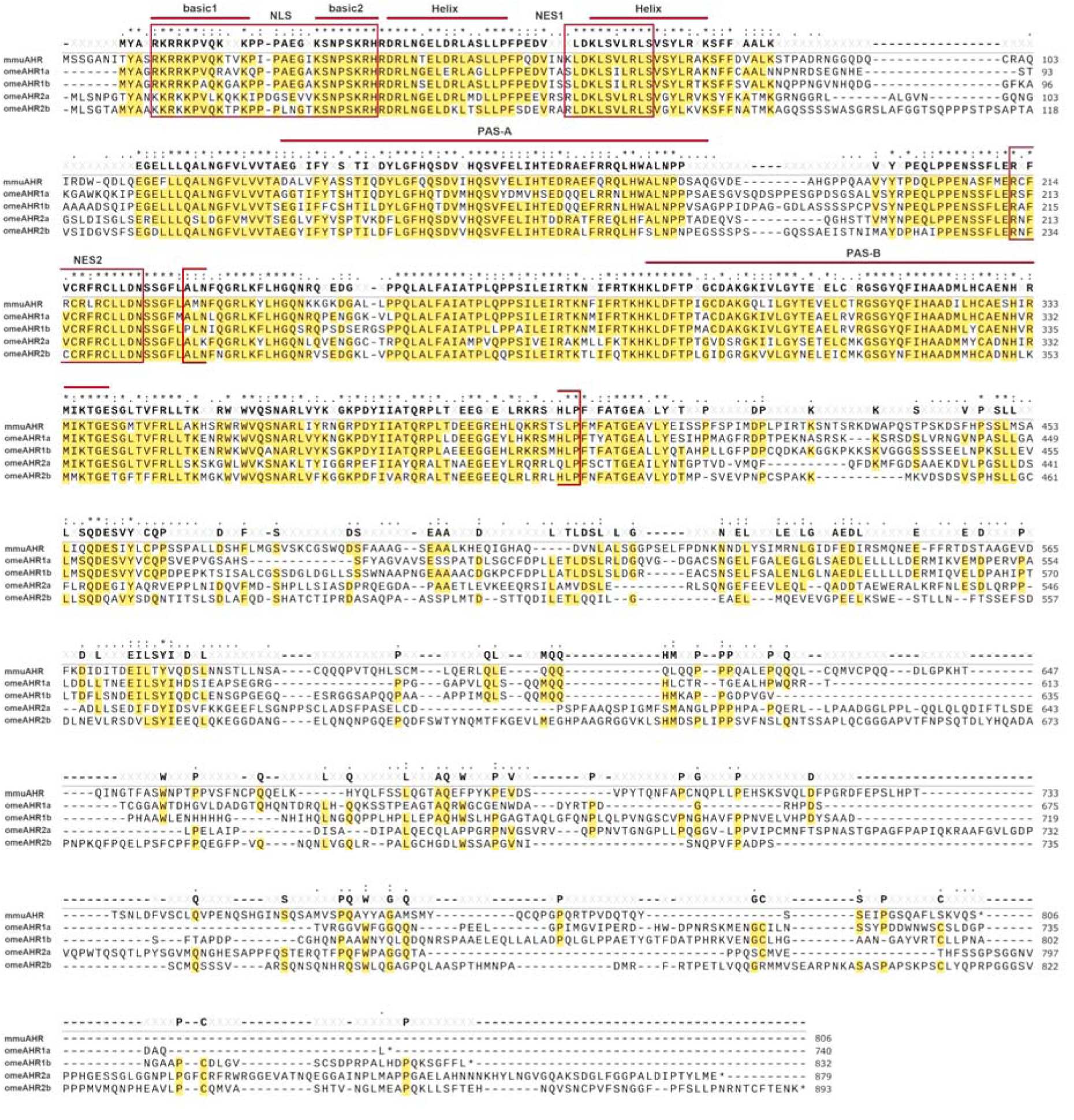
The alignment of AHRs using Clustal Omega, including omeAHR1a, omeAHR1b, omeAHR2a, omeAHR2b, and mmuAHR. Key functional domains were pinpointed by aligning them with the mouse AHR reference sequence ^46^, including NLS ^47^, NES1 ^47^, and NES2 ^48^, are highlighted within boxes for clarity. The LBD ^49^ is indicated by brackets. Identical sequences are emphasized through shading.

The phylogenetic analysis and alignment underscore the sequence diversity of AHRs across species. Our study also aims to explore the functional differences among AHR subforms or species, with a focus on subform-specific expression patterns and sensitivity to AHR ligands.

### 3.2 Marine medaka AHR and ARNT expression

The expression abundance of AHR is crucial given the tissue-and developmental stage-specific expression patterns that could influence physiological or toxicological responses. For example, zebrafish dreAHR2 (unlike dreAHR1a or 1b) plays a critical role in craniofacial and fin development and TCDD-dependent cardiotoxicity ^14, 29^. The gene expression levels of the AHR pathway are highly variable, suggesting that the predominance of certain forms should be investigated. In teleosts, AHR2 is often the predominant form, as seen in the Atlantic cod, where AHR2 is more abundant in the gill, liver, and brain ^30, 31^. However, in chickens, the AHR1-ARNT1 pair appears to be dominant in most tissues and is less sensitive to TCDD but sensitive to other compounds ^14^. This indicates that the differential expression of a dominant AHR or ARNT form can affect sensitivity and selectivity to specific compounds.

In marine medaka, omeAHR2a is more abundant than other AHR forms in embryo, gill, muscle, and brain (**Fig. 3**). OmeAHR2b generally has a greater abundance than omeAHR1a and omeAHR1b, except in the testis. For omeARNTs, the expression levels of omeARNT1 and omeARNT2 are comparable to that of the dominant omeAHR2a in the embryo and most tested tissues, similar to findings in common carp, where ccaARNT1 and ccaARNT2 are comparable to ccaAHR2 ^24^. In the brain and embryo, omeARNT2 is more abundant than omeARNT1. The pattern of greater ARNT2 abundance in the nervous system was also observed in other species like Atlantic cod and common carp, suggesting a conserved role for ARNT2. However, in other tested tissues, the expression levels of omeARNT1 and omeARNT2 are similar without significant difference. Considering omeAHRR as a significant partner and feedback mechanism for repressing AHR transactivation, its expression is generally much lower than that of omeAHR2a and omeARNT1/2 in all tested tissues. However, the expression of AHRR can be robustly induced in mouse liver or liver-derived cell lines of zebrafish upon exposure to potent AHR agonists, indicating the strong capability of AHR pathway itself in regulating the AHRR ^32, 33^.

**Figure 3.**
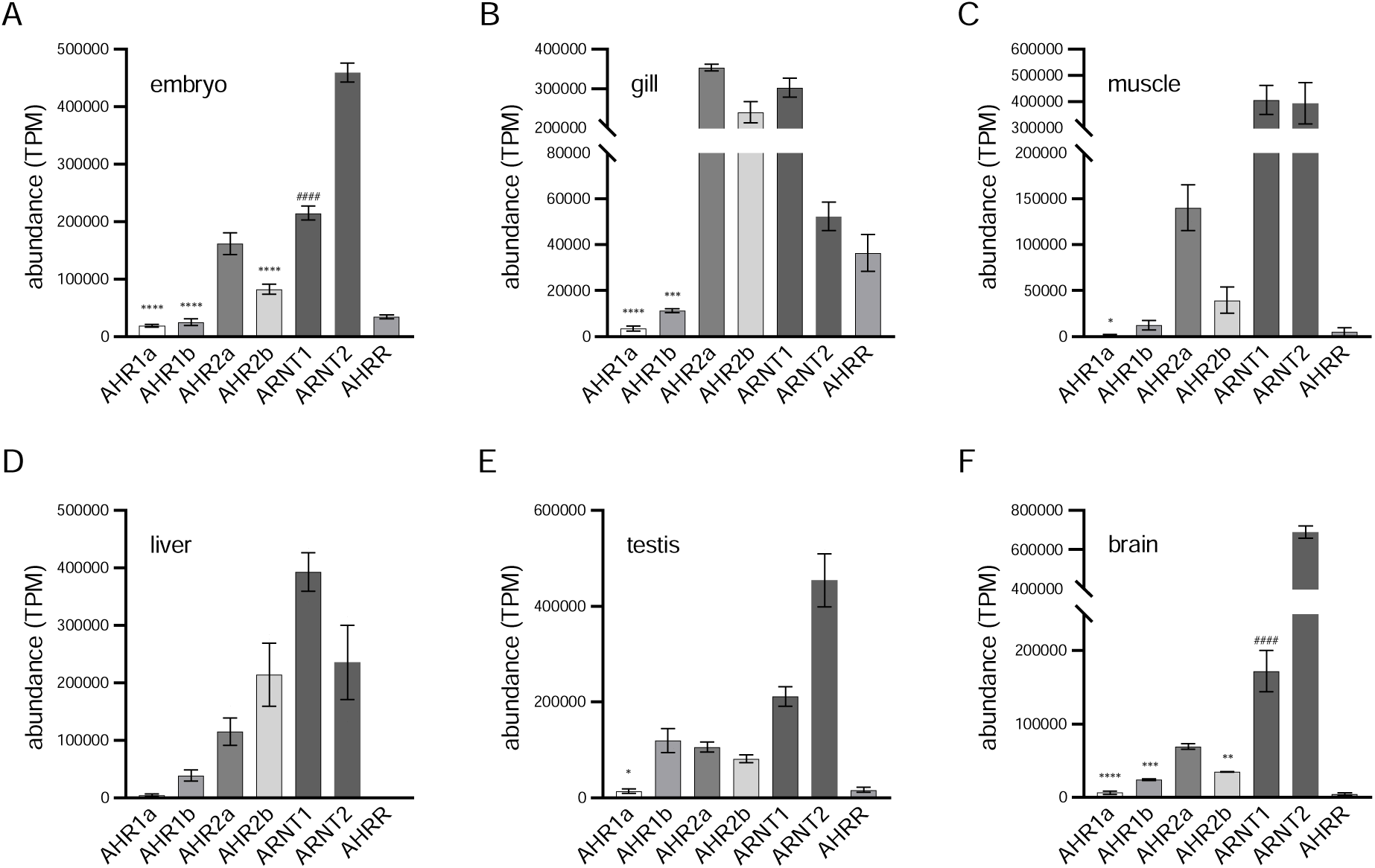
AHR pathway gene expression levels in the marine medaka embryo (A), gill (B), muscle (C), liver (D), testis (E), and brain (F). The expression levels are expressed in TPM (mean ± SD). Significant differences were tested by one-way ANOVA (or non-parametric test or t-test), and the results are presented as \**p* < 0.05, \*\**p* < 0.01, \*\*\**p* < 0.001, and \*\*\*\**p* < 0.0001 between AHRs (note: # marks the significance between ARNTs).

In various tissues of marine medaka, the expression patterns of all AHR pathway genes are similar, with omeAHR1a, omeAHR1b, and omeAHRR usually showing low abundances, while omeARNT1, omeARNT2, and omeAHR2a have comparable abundances. The omeAHR2a and omeAHR2b as the predominant forms, especially the omeAHR2a, potentially form the functional basis for physiological or toxicological outcomes.

### 3.3 Functional screening of omeAHR and omeARNT

In humans, as confirmed by in vitro studies using a reporter gene system, both ARNT and ARNT2 serve as partners to facilitate the function of AHR ^34^. However, ARNT2 exhibits only about 20% of the potency of ARNT. Comparative analysis of different ligands in activating AHR, such as TCDD, 3-methylcholanthrene, and benzo[a]pyrene, indicated that weak agonist preferentially induces the AHR-ARNT complex formation, whereas the potent ligand TCDD can efficiently induce dimer formation with both ARNT and ARNT2 ^35^. In the avian species chicken (*Gallus gallus*), the ggaAHR1-ggaARNT1 pair predominantly regulates the AHR response to natural ligands but not to TCDD. There is a nuanced functional difference between ggaARNT1 and ggaARNT2, with ggaAHR1 and ggaAHR2 being potently induced by TCDD in partnership with ggaARNT2 rather than ggaARNT1 ^36^. This background underscores the necessity for a detailed functional characterization of omeAHRs, including their cooperation potential with omeARNTs.

Using a luciferase reporter gene assay in COS-7 cells, the transactivation/cooperation potential of each omeAHR/omeARNT combinations under 100 nM TCDD exposure was measured. The results in **Fig. 4** show that, without omeARNTs (‘just AHR’ transfection mode), 3 out of the 4 omeAHRs were significantly induced by TCDD, including omeAHR1a (3.90× induction versus DMSO; **Fig. 4A**), omeAHR1b (2.21×; **Fig. 4B**), and omeAHR2b (2.87×; **Fig. 4D**). In the presence of omeARNTs (‘AHR + ARNT’ transfection mode), all omeAHRs exhibited robust transactivation in cooperation with omeARNTs. All omeAHRs cooperated well with omeARNTs, e.g., omeAHR1a-omeARNT1 (5.98× induction versus DMSO; **Fig. 4A**), omeAHR1b-omeARNT1 (5.03×; **Fig. 4B**), omeAHR2a-omeARNT1 (6.90×; **Fig. 4C**), and omeAHR2b-omeARNT1 (17.98×; **Fig. 4D**). OmeAHR1a/1b/2a cooperated with the omeARNT1/2 in similar degree of TCDD-induced transactivation, while omeAHR2b was weaker when partnered with omeARNT2 compared to omeARNT1. The situations seem not the same to Japanese medaka, such as certain combinations, olaAHR1a with olaARNT2 or dreARNT2, do not promote TCDD-induced transactivation. The data confirm the functionality of all omeAHRs and omeARNTs and set the stage for further exploration of their sensitivities.

**Figure 4.**
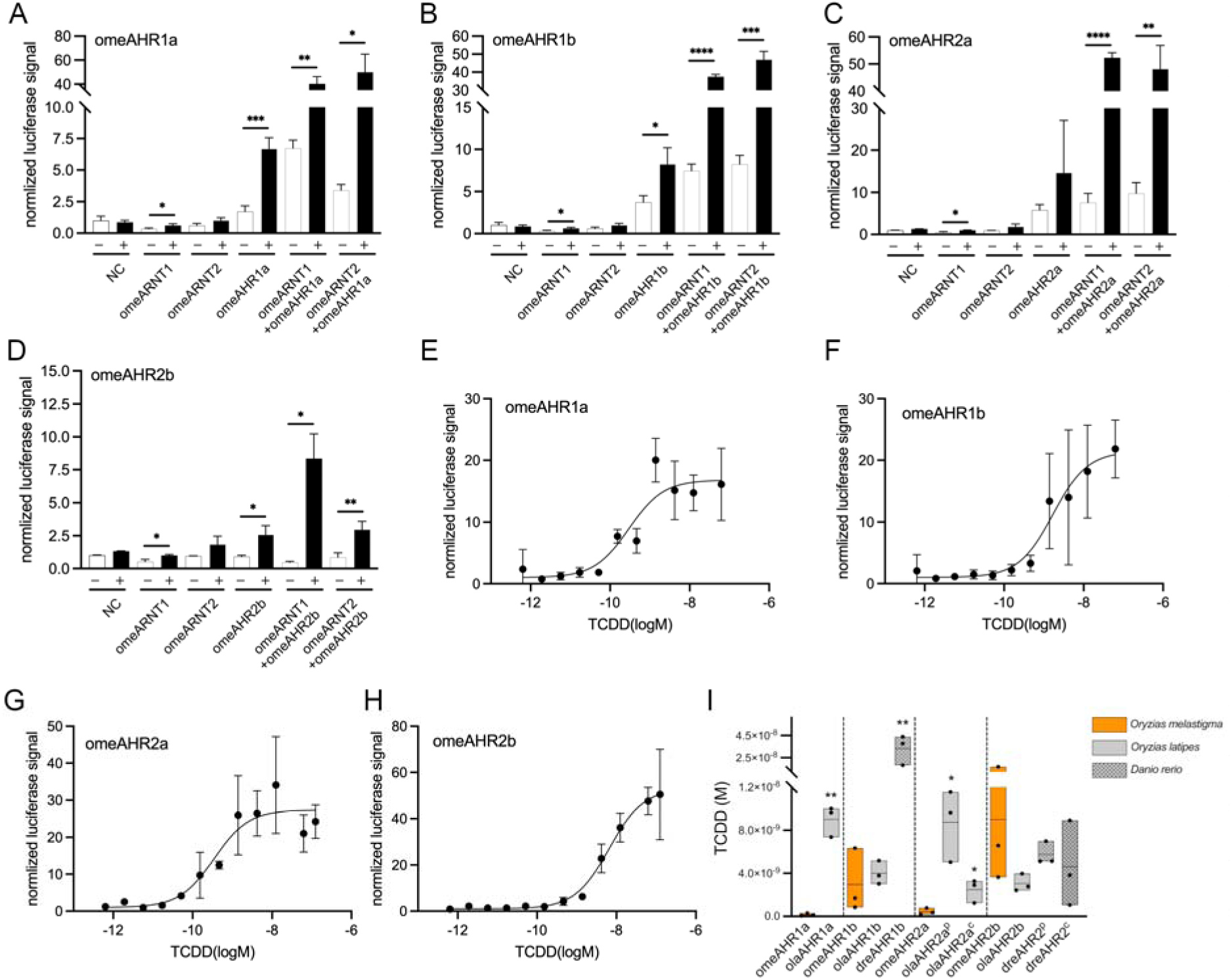
TCDD-induced (100 nM) transactivation of AHRs of marine medaka (omeAHR1a (A), omeAHR1b (B), omeAHR2a (C), and omeAHR2b (D)) with or without cotransfection with ARNT. The control (-) and TCDD (+) groups all have three biological replicates, and the date are expressed as the mean ± SD. Significant differences were tested by t-test. Dose-response curves of TCDD-induced AHR transactivation are: omeAHR1a (E), omeAHR2a (F), omeAHR1b (G), and omeAHR2b (H). Plot E shows the range of EC_50_ values, in which the ‘p’ mark on the gene name represents the data retrieved from our previous study ^23^ and the ‘c’ mark represents that determined in the current study. Significance was declared at the \**p* < 0.05, \*\**p* < 0.01, \*\*\**p* < 0.001, and \*\*\*\**p* < 0.0001.

### 3.5 Marine medaka’s in vivo/vitro sensitivity to TCDD

As an important toxicity testing model, marine medaka’s in vivo/vitro sensitivity to dioxin have not been so clearly characterized like the close relative Japanese medaka. Careful determination of the marine medaka’s response to dioxin/DLCs could enable the precise toxicity evaluation of pollutants particularly for the marine environment. Previous reports on Japanese medaka have examined the toxicity of TCDD, determining the LC_50_ values to be 27.1 pM (equivalent to 8.72 ng/L at 21 days post-fertilization) ^37^ and 13.5 ng/L (at 3 days post-hatching) ^38^, with corresponding EC_50_ values for TCDD-induced hatching failure of 47.1 ng/L ^37^ and 26.8 ng/L ^38^, respectively. For a more accurate evaluation and consistency in comparison, we also determined TCDD-induced mortality and hatching failure using both marine and Japanese medaka (**Fig. S1**). In Japanese medaka, similar to previously reports, the LC_50_ at 3 days post-hatching is 3.42 ng/L (95% CI: 1.37-6.48 ng/L; **Fig. 5B**), and the EC_50_ of hatching failure is 50.16 ng/L (**Fig. S1B**). In comparison, in marine medaka the LC_50_ and EC_50_ are 1.64 ng/L (95% CI: 1.05-2.55 ng/L; **Fig. 5A**) and 35.67 ng/L (**Fig. S1D**), respectively. The above data provide evidence at the physiological level that the marine medaka is more sensitive to TCDD than the Japanese medaka. However, the underlying mechanisms remain unknown, and whether these differences are mediated by the AHR will be explored below at the in vitro and in silico molecular levels.

**Figure 5.**
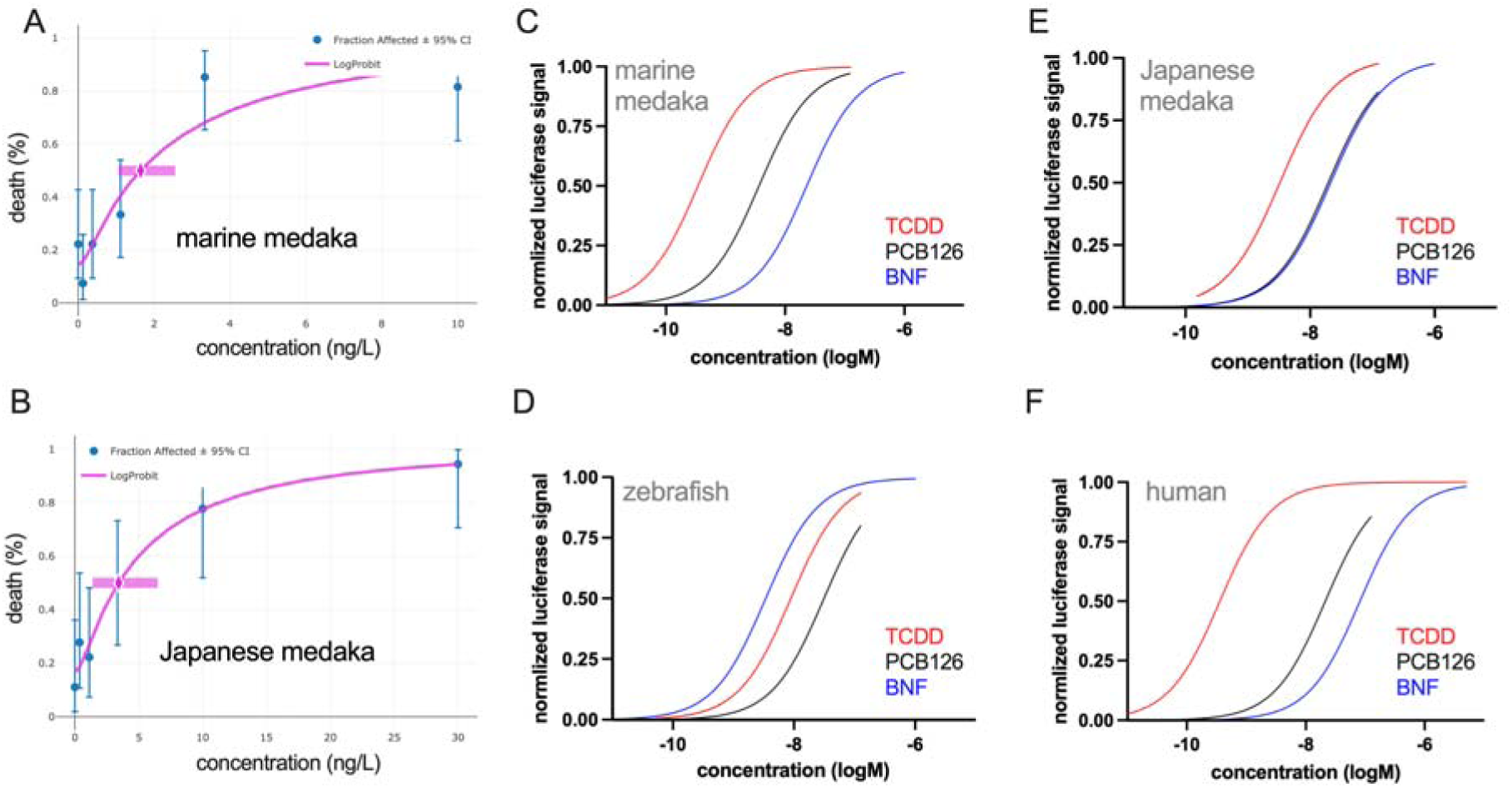
Embryo toxicity of TCDD and species-specific in vitro AHR activation potency of additional AHR ligands in marine/Japanese medaka. (A) and (B) are dose-response curves of TCDD-induced embryo death at the time-point of ‘3-day post hatching’ with LC_50_ and 95% CI indicated. (C-F) Transactivation curves of omeAHR2a, dreAHR2, olaAHR2a, and hsaAHR, respectively.

The species-specific toxicity of compounds is a cornerstone of ecotoxicology evaluation, with the AHR being a key mediator of the toxicity of dioxins/DLCs. The in vitro assessment of AHR sensitivity in different species can predict potential toxic outcomes in a given species, making it a valuable tool for ecotoxicological research ^13, 39^. Marine medaka, a widely used model species, has shown alterations in gene expression within the AHR pathway due to exposure to dioxins and DLCs. However, a systematic determination of the sensitivity and functionality of the marine medaka AHR pathway has been lacking, which this study aims to address. The study compared the sensitivities of marine medaka, Japanese medaka, zebrafish, and human AHRs to the prototypical dioxin congener—TCDD as well as additional ligands.

As reported by our previous research, there are four AHRs in Japanese medaka, and the four AHR subforms’ sensitivities to TCDD were similar and all located in the nM range, e.g., the EC_50_ values of olaAHR1a, 1b, 2a, and 2b are respectively 9.01±1.43 nM, 4.00±1.10 nM, 8.75±3.34 nM, and 3.06±0.81 nM ^23^. While in marine medaka, we determined in this study, omeAHR1a exhibited clear TCDD-induced transactivation when co-transfected with omeARNT1, with an EC_50_ of 0.16±0.12 nM (**Fig. 4E**). The relative sensitivity (ReS) of omeAHR1a is 36 (**Table 1**), indicating a greater sensitivity compared to Japanese medaka olaAHR1a (ReS = 0.64) and zebrafish dreAHR2- dreARNT1 (ReS = 1.00; serves as the standard for parallel comparison across AHR subforms). As the dominant form, omeAHR2a also shows greater sensitivity, with an EC_50_ of 0.44±0.30 nM (**Fig. 4G**) and ReS of 13.09 (**Table 1**). The EC_50_ values for omeAHR1b (**Fig. 4F**) and omeAHR2b (**Fig. 4H**) are 2.96 ± 2.96 nM and 9.00 ± 6.88 nM, respectively, falling within a similar range to that of dreAHR2 (**Table 1**). In comparison with zebrafish, all omeAHRs and dreAHR2 are much sensitive than the dreAHR1b (EC_50_: 33.30±13.37 nM) ^23^. However, as the dominant form in zebrafish, dreAHR2 maintains an EC_50_ at the nM level that is comparable to other fish species’ dominant AHR subforms. When compared with hsaAHR, the highly sensitive subforms (omeAHR1a and omeAHR2a) appear to be in the same range as hsaAHR (EC_50_ = 0.43±0.30 nM, **Table 1**). Collectively, by performing a subform-based comparison (**Fig. 4I**), omeAHR1a was found to be statistically more sensitive than olaAHR1a. Both omeAHR1b and olaAHR1b are more sensitive than dreAHR1b. OmeAHR2a is more sensitive than olaAHR2a. No significant differences were observed among omeAHR2b, olaAHR2b, and dreAHR2 (note that: dreAHR2 belongs to the AHR2b subfamily according to the phylogenetic analysis in **Fig. 1A**).

**Table 1.** TCDD-induced EC_20/50/80_ values of AHR activation and the relative sensitivity (ReS) of AHR.

| AHR | EC<sub>20</sub> (nM) | EC<sub>50</sub> (nM) | EC<sub>80</sub> (nM) | Top/Bottom | ReS* | References |
| --- | --- | --- | --- | --- | --- | --- |
| omeAHR1a | 0.04±0.03 | 0.16±0.12 | 0.62±0.49 | 11.47±4.66 | 36.00 | current study |
| omeAHR1b | 0.74±0.74 | 2.96±2.96 | 11.82±11.84 | 49.88±57.16 | 1.95 | current study |
| omeAHR2a | 0.11±0.08 | 0.44±0.30 | 1.75±1.22 | 25.92±10.28 | 13.09 | current study |
| omeAHR2b | 2.25±1.72 | 9.00±6.88 | 35.98±27.50 | 45.91±6.85 | 0.64 | current study |
| olaAHR1a | 2.25±0.36 | 9.01±1.43 | 36.03±5.75 | 3.45±0.33 | 0.64 | <sup>23</sup> |
| olaAHR1b | 1.00±0.27 | 4.00±1.10 | 15.99±4.39 | 4.19±1.00 | 1.44 | <sup>23</sup> |
| olaAHR2a | 2.19±0.84 | 8.75±3.34 | 35.01±13.36 | 6.59±0.35 | 0.66 | <sup>23</sup> |
| olaAHR2a | 0.62±0.27 | 2.48±1.08 | 9.91±4.33 | 11.24±3.63 | 2.32 | current study |
| olaAHR2b | 0.76±0.20 | 3.06±0.81 | 12.22±3.25 | 6.90±0.49 | 1.88 | <sup>23</sup> |
| dreAHR1b | 8.88±3.45 | 33.30±13.37 | 133.32±53.59 | 3.38±0.27 | 0.17 | <sup>23</sup> |
| dreAHR2 | 1.44±0.27 | 5.76±1.08 | 23.01±4.28 | 4.41±0.62 | 1.00* | <sup>23</sup> |
| dreAHR2 | 1.15±1.00 | 4.60±3.99 | 18.39±15.94 | 3.26±1.20 | 1.00* | current study |
| hsaAHR | 0.11±0.08 | 0.43±0.30 | 1.71±1.21 | 91.77±117.02 | 13.40 | current study |

\* ReS = (EC<sub>50</sub> of dreAHR2)/(EC<sub>50</sub> of interest)<sup>45</sup>, the EC<sub>50</sub> of dreAHR2 = 5.76±1.08 nM<sup>23</sup> was used for ReS calculation. Each AHR cooperates with the ARNT1 subform of the corresponding species, e.g., omeAHR1a-omeARNT1 and dreAHR2-dreARNT1.

Considering additional reported fish AHRs, lake sturgeon and white sturgeon AHRs exhibit higher sensitivity to TCDD ^40^, whereas the Atlantic cod AHR2a is less sensitive, with an EC_50_ greater than 10 nM ^41^. Collectively, both the in vivo and in vitro data show good agreement and support the conclusion that marine medaka is more sensitive than its close freshwater relative, the Japanese medaka. This supports the use of marine medaka as a potentially highly sensitive model for toxicity evaluation, at least for dioxins in marine environments. However, the above findings also underscore the species-and subform-specific sensitivities to TCDD, indicating that caution should be exercised when using data from one species to predict the potential outcomes in another species.

### 3.6 The sensitivity of AHR to additional ligands among species

To gain a more comprehensive understanding of the functions of omeAHRs and other AHRs, as well as their species-specific properties, we determined the responsiveness of these receptors (the predominant forms of each species: omeAHR2a, olaAHR2a, dreAHR2, and hsaAHR) to additional ligands, including PCB126, BNF, and indole. All chemicals were first tested by MTT assay to ensure the exposure concentration range do not seriously alter the cell viability (**Fig. S2** and **S3** for COS-7 and HepG2 cells, respectively). PCB126 is a dioxin-like compound that usually produces similar toxicity to traditional dioxin. In comparison with TCDD, the PCB126-induced EC_50_ values for the transactivation of omeAHR2a, olaAHR2a, dreAHR2, and hsaAHR are 5.52±1.64 nM, 10.49±7.80 nM, 62.39±30.57 nM, and 13.74±6.89 nM, respectively (**Fig. 5C/5D/5E/5F** and **Table 2**). The average EC_50_ for PCB126-induced transactivation across the four AHRs (23.04 nM) is approximately ten times that of the TCDD-induced EC_50_ (1.99 nM). This finding aligns closely with the WHO-reported TEF of 0.1 for PCB126, compared to 1 for TCDD ^16^. As the dioxin-like compound, the EC_50_ trend produced by PCB126 (omeAHR2a < olaAHR2a < hsaAHR < dreAHR2) is quite resemble that by TCDD (hsaAHR ≈ omeAHR2a < olaAHR2a < dreAHR2). The dreAHR2, most insensitive to TCDD also the most insensitive to PCB126; and the most sensitive one is almost the omeAHR2a. It is evident that the data acquired from TCDD can be predictive for the same class of compounds, such as DLCs, particularly for PCB126 in this case. However, for other classes of compounds such as the BNF, the situation does not seem to follow the same pattern as indicated by TCDD.

**Table 2.** EC_50_ of omeAHR2a, olaAHR2a, dreAHR2, and hsaAHR under the TCDD, PCB126, and BNF activation. The data were expressed as mean ± SD in nM units.

| **AHR** | **EC<sub>50</sub>-TCDD** | **EC<sub>50</sub>-PCB126** | **EC<sub>50</sub>-BNF** | **ReP<sub>PCB126</sub>** | **ReP<sub>BNF</sub>** |
| --- | --- | --- | --- | --- | --- |
| omeAHR2a | 0.44±0.30 | 5.52±1.64 | 46.38±20.07 | 0.08 | 0.01 |
| olaAHR2a | 2.48±1.08 | 10.49±7.80 | 19.36±1.98 | 0.24 | 0.07 |
| dreAHR2 | 4.60±3.99 | 62.39±30.57 | 2.12±1.19 | 0.07 | 2.17 |
| hsaAHR | 0.43±0.30 | 13.74±6.89 | 116.2±44.17 | 0.03 | 0.0037 |

BNF belongs to a class of flavonoids, and is able to activate AHR and induce phase I and phase II metabolic enzymes involved in the metabolism and detoxification of xenobiotics. Flavonoids typically exist in plants as secondary metabolites and have various biological activities such as antioxidant, anti-inflammatory, and anticancer properties. The order for the EC_50_ values of BNF in transactivating AHR is dreAHR2 < olaAHR2a < omeAHR2a < hsaAHR (**Fig. 5C/5D/5E/5F** and **Table 2**). It is clear that the AHR form least sensitive to TCDD, dreAHR2, has become the most sensitive to BNF. And interestingly, the most sensitive form in transactivation, hsaAHR, becomes the least sensitive to BNF. Therefore, the insights gained from prototypical dioxins cannot be directly applied to other types of AHR ligands.

Based on the classification of AHR ligands into full agonists, partial agonists, and antagonists, the above data indicate that TCDD, PCB126, and BNF act as potent full agonists. The segment of partial or weak agonists has not been tested and is explored here. Indole itself, as well as other indole derivatives produced by gut microbiota from dietary tryptophan, can act as endogenous ligands for AHR. Previous reports have determined the indole has a EC_50_ ∼ 3 μM in the luciferase reporter gene assays ^42^. In addition, some naturally occurring marine brominated indoles serve as AHR ligands/agonists. However, possibly due to indole’s rapid metabolism, the exposure time in the luciferase reporter gene assay was slightly modified to use an 8-hour exposure period ^43^. However, the current study did not obtain significant activation in the transactivation assay. Possible reasons may include the use of the COS-7 cell line or the current system not being sensitive enough to detect weak agonists like indole.

### 3.7 In silico modeling the interaction between omeAHRs-DLCs

In silico modeling, including virtual docking and molecular dynamics (MD) analysis, was conducted to explore the interaction between AHRs and ligands. Such analysis may shed light on explaining the molecular mechanism of sensitivity differences between marine medaka and Japanese medaka, as well as zebrafish and human. Initial analysis identified similar binding cavity volumes within the LBD of the four AHRs: omeAHR2a (117.190 Å^3^), olaAHR2a (117.100 Å^3^), dreAHR2 (121.240 Å^3^), and hsaAHR (102.840 Å^3^). Docking studies corroborated transactivation data, demonstrating that TCDD and PCB126 can effectively bind to various AHR forms with similar conformations, e.g., by interacting with the HIS-284, PHE-288, PHE-344, ILE-318 and LEU-346 of omeAHR2a (**Fig. 6A and 6B**), similarly, residues confer the interaction with olaAHR2a (**Fig. 6E and 6F**) are the same (e.g., HIS-277, PHE-281, PHE-337, ILE-311, and LEU- 339). In detail, the omeAHR2a interacts with TCDD through hydrophobic interaction (ILE-318) and π-stacking (HIS-284, PHE-288 and PHE-344) with TCDD. The interactions of omeAHR2a- BNF (**Fig. 6C**) become more complex in compare with omeAHR2a-TCDD, which involve additional residues conferring hydrophobic interactions (ASN-329 and PRO-290), and extra hydrogen bond with SER-358. Given that indole has a smaller chemical spatial structure, its conformation when binding with the omeAHR2a is more flexible (**Fig. 6D** and **6L**), and through specific amino acid residues (HIS-300) different from TCDD. As a whole, the omeAHR2a and olaAHR2a have much greater sequence similarity, and bind with the ligands through almost the same poses, while in contrast much different from which of hsaAHR and dreAHR2 (**Fig. S4**).

**Figure 6.**
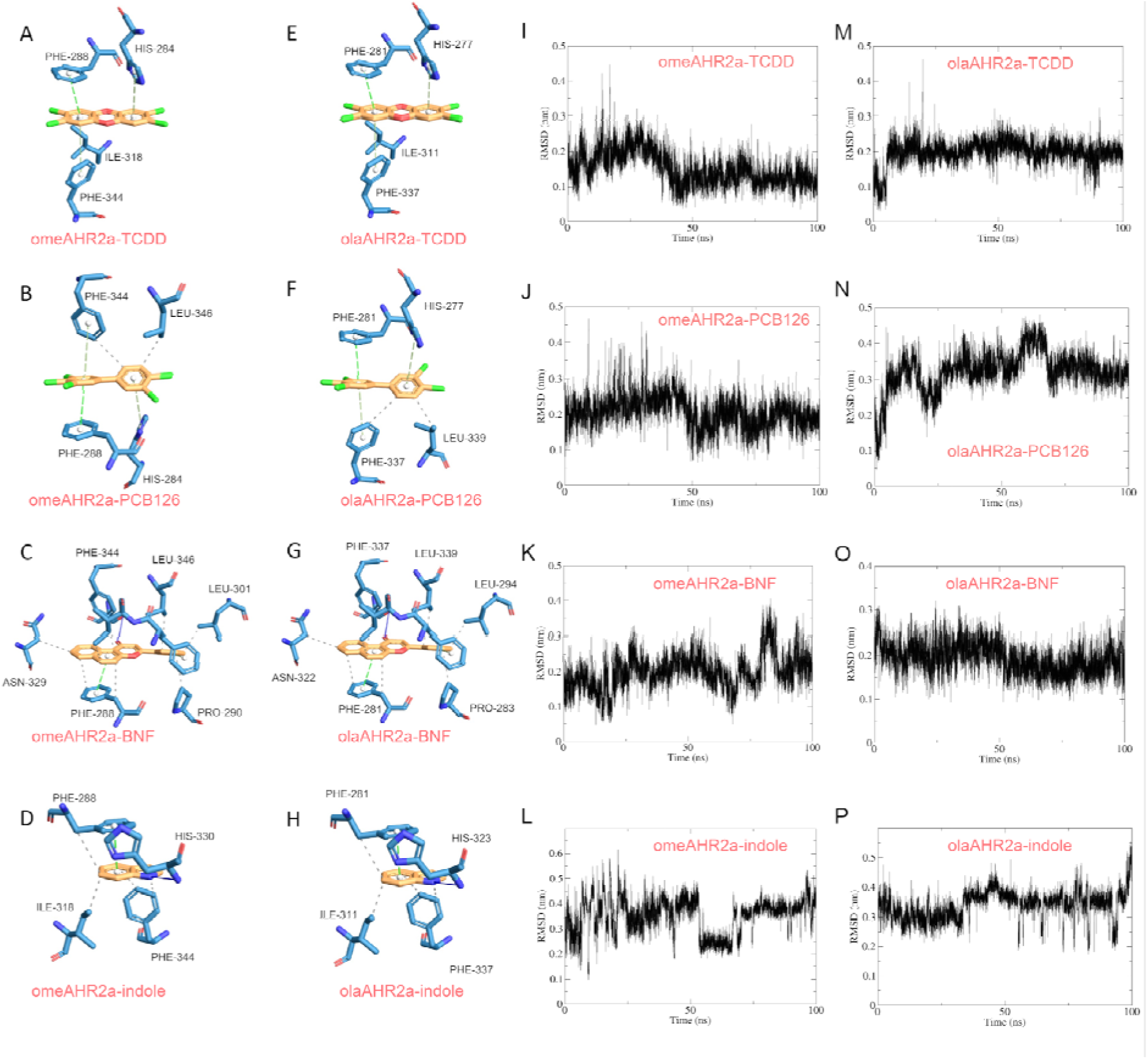
Molecular docking modes for AHR-ligand interaction and the ligands’ RMSD (0-100 ns), including A and I (omeAHR2a-TCDD), B and J (omeAHR2a-PCB126), C and K (omeAHR2a- BNF), D and L (omeAHR2a-indole), E and M (olaAHR2a-TCDD), F and N (olaAHR2a-PCB126), G and O (olaAHR2a-BNF), and H and P (olaAHR2a-indole).

After the molecular dynamics analysis (**Fig. 6**, **S5**, and **S6**), we’ve found that even omeAHR2a and olaAHR2a have similar binding poses, the dynamics of their interactions with ligands are more different (**Fig. 6**). The omeAHR2a interacts with TCDD in a more stable conformation with the RMSD mostly below 0.2 nm (**Fig. 6I**) while the olaAHR2a always slightly greater than 0.2 nm (**Fig. 6M**). For PCB126, the situations are similar that omeAHR2a interacted with PCB126 more stably and had less RMSD (around 0.2 nm; **Fig. 6J**). On the contrary, the olaAHR2a interacted with BNF in a more stable dynamics state in comparison with omeAHR2a. When interacting with indole, each AHR subform indicated the fluctuate of RMSD (**Fig. 6L, 6P, S5**, and **S6**) which means the unstable state of the interactions and possible less binding affinity. Finally, through the MM-PBSA calculation, the binding affinity data (**Table 3**) actually confirm the above phenomenon that, omeAHR2a-TCDD/PCB126 interactions are more stable with less △G of-30.75/-26.82 kcal/mol, which indicates much stable binding than olaAHR2a-TCDD/PCB126 (△G of-26.88/-22.16 kcal/mol). This seems that, at least for dioxin and DLCs, the omeAHR2a has greater binding affinity and less dynamics flexibility than olaAHR2a, and which may ensure the greater sensitivity in omeAHR2a. In contrast, the △G of olaAHR2a-BNF (-29.95 kcal/mol) is less than that of omeAHR2a (-24.17 kcal/mol). In compare with other ligands, indole has much greater △G (∼-10 kcal/mol) in interacting with any of the tested AHR forms (**Table 3**).

**Table 3.** Binding affinity and binding cavity volume before and post ligand-binding.

| binding affinity and volume | omeAHR2a | olaAHR2a | hsaAHR | dreAHR2 |
| --- | --- | --- | --- | --- |
| binding energy-TCDD ($\Delta G$, kcal/mol) | -30.751 | -26.884 | -30.204 | -30.897 |
| binding energy-PCB126 ($\Delta G$, kcal/mol) | -26.821 | -22.162 | -24.137 | -28.463 |
| binding energy-BNF ($\Delta G$, kcal/mol) | -24.175 | -29.951 | -23.072 | -26.866 |
| binding energy-indole ($\Delta G$, kcal/mol) | -10.106 | -8.932 | -11.835 | -9.993 |
| cavity volume (initial; $\text{\AA}^3$) | 117.190 | 117.100 | 102.840 | 121.240 |
| cavity volume after MD-TCDD ($\text{\AA}^3$) | 166.777 | 186.199 | 196.088 | 203.585 |
| cavity volume after MD-PCB126 ($\text{\AA}^3$) | 357.376 | 170.237 | 251.561 | 265.670 |
| cavity volume after MD-BNF ($\text{\AA}^3$) | 171.408 | 204.283 | 280.395 | 166.572 |
| cavity volume after MD-indole ($\text{\AA}^3$) | 188.928 | 146.703 | 127.769 | 150.643 |

As a whole, we’ve found that among the tested ligands, the binding affinity usually follows this order: TCDD > PCB126 > BNF > indole (**Table 3**). The TCDD usually induces the various AHRs binding cavity to accommodate into a relatively compact volume larger than that of PCB126, while that might be bigger than that of indole due to its relatively smaller molecular space occupation (**Table 3**). It seems that, in most cases, in a given AHR subform, the greater binding affinity may produce increased AHR sensitivity to ligands. But this rule would not always work well across distinct AHR subforms, e.g., similar binding affinity (or even greater) does not ensure a more sensitive AHR phenotype in reacting with a certain ligand, e.g., when binding with TCDD, the dreAHR2 has greater binding affinity than omeAHR2a but less sensitivity indicated by EC_50_. And the tested ligands always obey a rule that their EC_50_ follow the order of TCDD < PCB126 < BNF, and indole does not produce a clear transactivation curve under current testing method. And even distinct AHRs can have differential sensitivity, the ranking order of AHR-transactivation potency of ligands may not change greatly (**Fig. 5 and Table 2**). Moreover, obvious species-specific and subform-specific phenomenon can also occur, such as the BNF (2.12±1.19 nM) can significantly activate the dreAHR2 with a much less EC_50_ than TCDD (4.60±3.99 nM, **Table 2**). Such situation also happens in a zebrafish hepatocytes cells ZFL transfected with pGL4.43L+LpRL-null reporter constructs, and produce similar trend that BNF (490 pM) owns less EC_50_ than TCDD (2.70 nM) ^44^.

## 4. Conclusion

This study provides a comprehensive analysis of the in vivo/vitro difference between two key species for dioxin ecotoxicological testing, marine medaka (*Oryzias melastigma*) and its close relative —Japanese madaka (*Oryzias latipes*), focusing on their sensitivity to dioxins and DLCs. The results indicate the in vitro sensitivity of AHR among species can vary by one or two orders of magnitude, with marine medaka showing heightened sensitivity to TCDD compared to Japanese medaka. This is further supported by in vivo toxicity testing, which aligned with the in vitro findings. The study also investigated the interaction of AHRs of various species with additional ligands, revealing that while TCDD and PCB126 follow a predictable pattern of transactivation, other compounds like BNF and indole do not adhere to the same sensitivity profile. In silico modeling of the AHR-ligand interaction provides insights into the molecular basis of the observed sensitivities, omeAHR2a, olaAHR2a, dreAHR2, and hsaAHR interact with ligands mostly in the affinity order of TCDD > PCB126 > BNF > indole, mirroring their AHR transactivation potency, but the docking poses and dynamics can vary. Overall, the study underscores the necessity for nuanced risk assessments that consider species-specific and subform-specific AHR sensitivities when evaluating the toxicity of environmental pollutants, and contributes valuable data for the accurate prediction of toxic effects in marine environments and advances the application of marine medaka as a model species in ecotoxicology.

## ACKNOWLEDGEMENTS

This work was supported by the National Natural Science Foundation of China (No. 22006049) and the Key Research and Development Project of Jining City (No. 2021JNZC076).

All the authors wrote and revised this paper. The authors declare no competing financial interests.

