## Supplemental Material for "Marine medaka responds differently to dioxin compared with its close freshwater relative, the Japanese medaka: the AHR molecular mechanism"

**Table S1**. NCBI accessions of the sequences involved in the phylogenetic analysis.

| Species | NCBI Accession (except for olaARNT2) |
| --- | --- |
| *Oryzias latipes* | olaAHR1a (XM_011489981.3), olaAHR1b (NM_001104678.1), olaAHR2a (XM_023950680.1), olaAHR2b (XM_011481447.3), olaARNT1 (XM_020710236.2), and olaARNT2 (Ensembl Accession: ENSORLP00015010476.1) |
| *Danio rerio* | dreAHR1a (NM_131028.2), dreAHR1b (NM_001024816.2), dreAHR2 (NM_131264.1), dreARNT1 (XM_005157852.4), and dreARNT2 (NM_131674.1) |
| *Takifugu rubripes* | truAHR1a (NM_001037962.1), truAHR1b (NM_001037959.1), truAHR2a (NM_001037960.1), truAHR2b (NM_001037963.1), and truAHR2c (NM_001037958.1) |
| *Anabas testudineus* | ateAHR1a (XM_026376424.1), ateAHR1b (XM_026361325.1), ateAHR2a (XM_026375976.1), ateAHR2b (XM_026363930.1), ateARNT1 (XM_026373162.1), and ateARNT2 (XM_026378163.1) |
| *Cyprinus carpio* | ccaAHR1a (XM_042740816.1), ccaAHR1b (XM_042748578.1), ccaAHR2 (XM_042749540.1), ccaARNT1 (XM_042771954.1), and ccaARNT2 (XM_042759473.1) |
| *Caenorhabditis elegans* | celAHR (NM_001025865.4) and celARNT (NM_001264398.3) |
| *Rattus norvegicus* | rnoARNT1 (NM_012780.4) and rnoARNT2 (NM_012781.4) |
| *Homo sapiens* | hsaARNT (NM_001668.4) and hsaARNT2 (NM_014862.4) |
| *Mus musculus* | mmuARNT (NM_001037737.2) and mmuARNT2 (NM_007488.4) |

**Table S2.** Primers for marine medaka AHR and ARNT cloning.

| **Gene** | **Primer sequence (5′→ 3′)** |
| --- | --- |
| omeAHR1a | Forward: AGAAGATTCTAGAGCTAGCGATGTACGCGGGACGGAAGAGGAGGA  Reverse: ATCCGATTTAAATTCGAATTTCAGAGCTGAGCGTCCGGACCGTCC |
| omeAHR1b | Forward: AGAAGATTCTAGAGCTAGCGATGTACGCCGGGCGCAAACGCAGGA  Reverse: ATCCGATTTAAATTCGAATTTCAAAGAAAGAATCCGGATTTCTGG |
| omeAHR2a | Forward: AGAAGATTCTAGAGCTAGCGATGTTGAGTAACCCGGGAACGTACG  Reverse: ATCCGATTTAAATTCGAATTCTACTCCATCAGGTACGTGGGGATG |
| omeAHR2b | Forward: AGAAGATTCTAGAGCTAGCGATGCTGTCCGGCACCGCCATGTACG  Reverse: ATCCGATTTAAATTCGAATTCTACTTGTTCTCGGTGAAGCAGGTG |
| omeARNT1 | Forward: AGAAGATTCTAGAGCTAGCGATGTTATTTCACACAGACATGACCT  Reverse: ATCCGATTTAAATTCGAATTTCATTCATTGTATTGAGGGTAGAGG |
| omeARNT2 | Forward: AGAAGATTCTAGAGCTAGCGATGGCAACCCCCGCCGCGGTCAACC  Reverse: ATCCGATTTAAATTCGAATTCTACTCCGAGAAGTTGGGGAACATG |

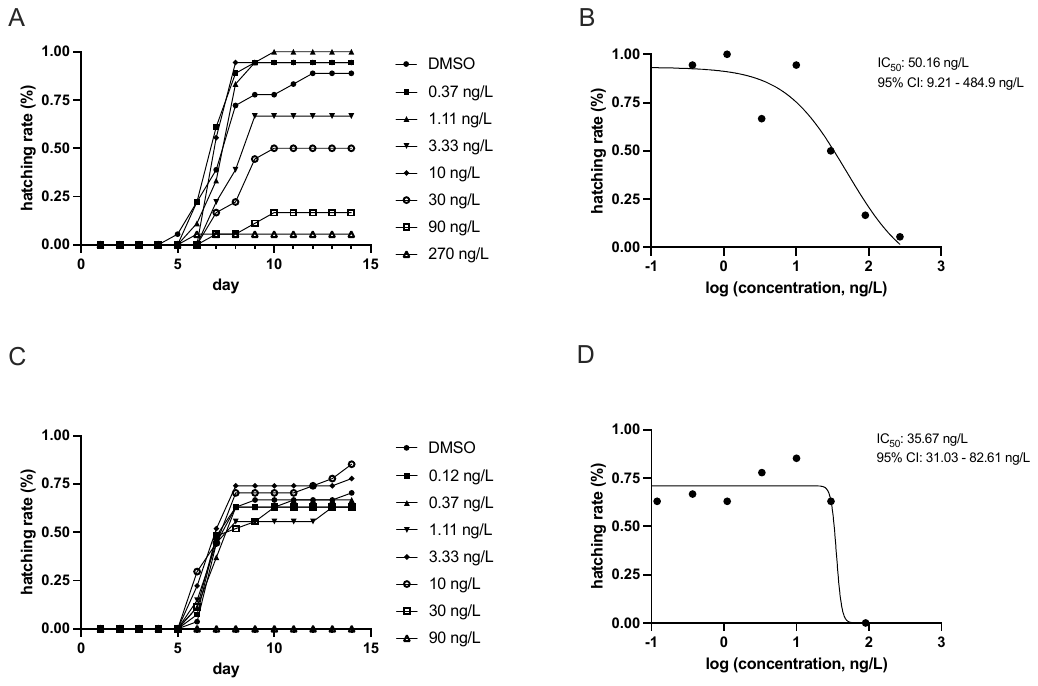

**Figure S1.** Hatching rate of marine medaka and Japanese medaka under TCDD exposure. (A) The time-course of Japanese medaka hatching rate under various concentrations of TCDD exposure. (B) The IC_50_ of Japanese medaka’s hatching rate under TCDD exposure at day 13 after exposure. (C) The time-course of marine medaka hatching rate under various concentrations of TCDD exposure. (D) The IC_50_ of marine medaka’s hatching rate under TCDD exposure.

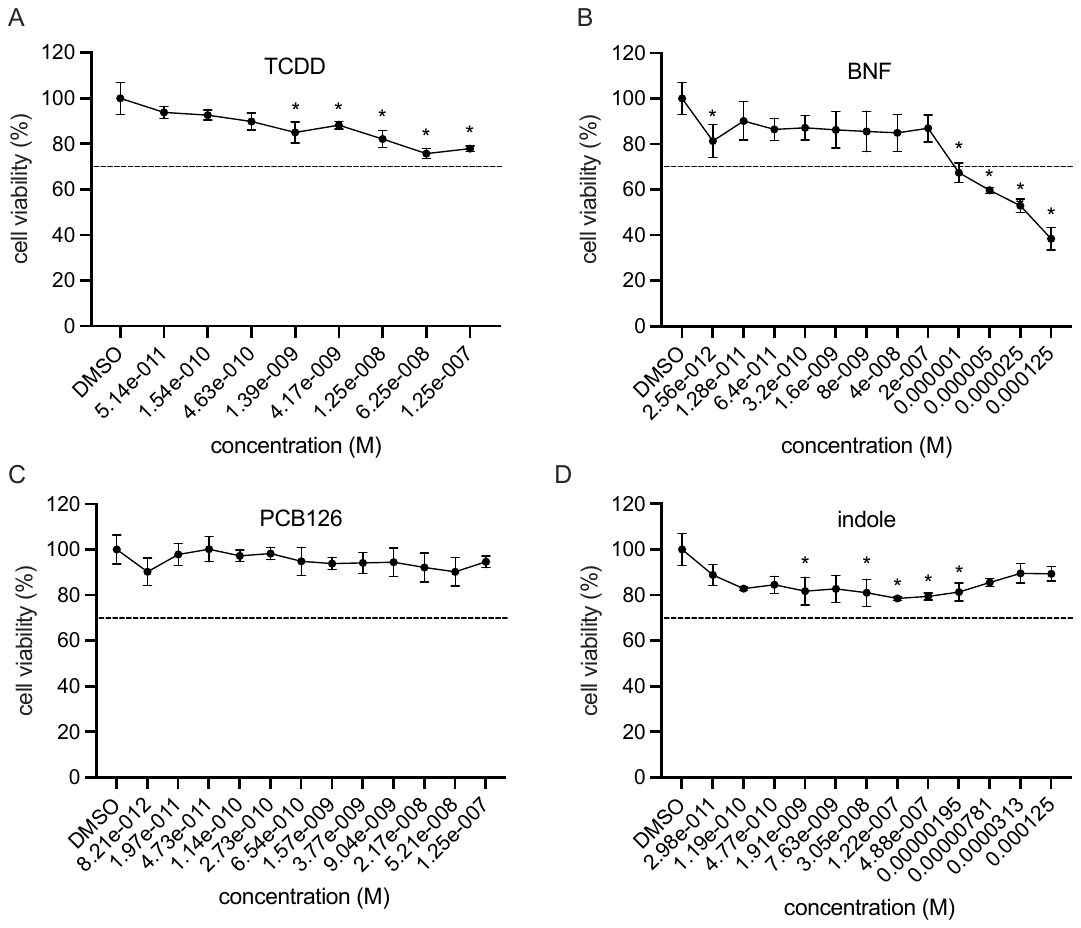

**Figure S2.** MTT results of TCDD, PCB126, BNF, and indole in COS-7 cells.

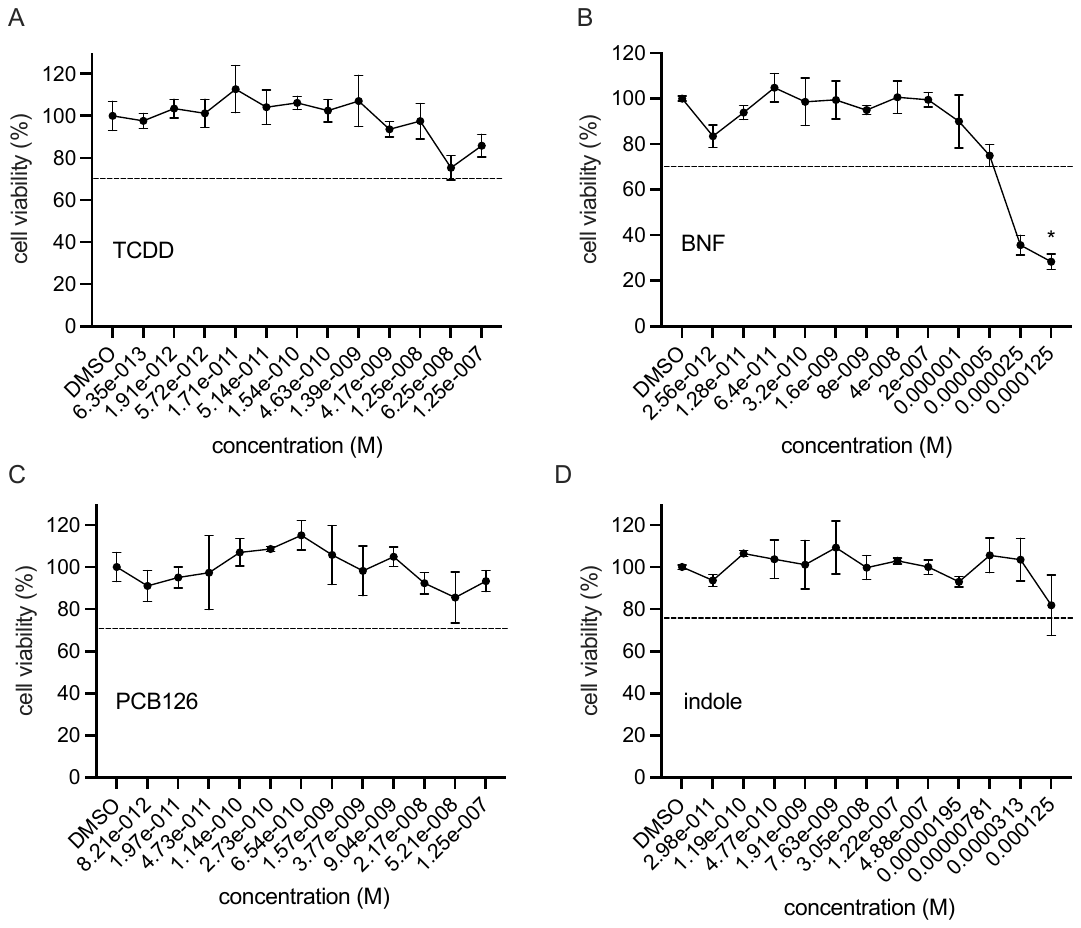

**Figure S3.** MTT results of TCDD, PCB126, BNF, and indole in HepG2 cells.

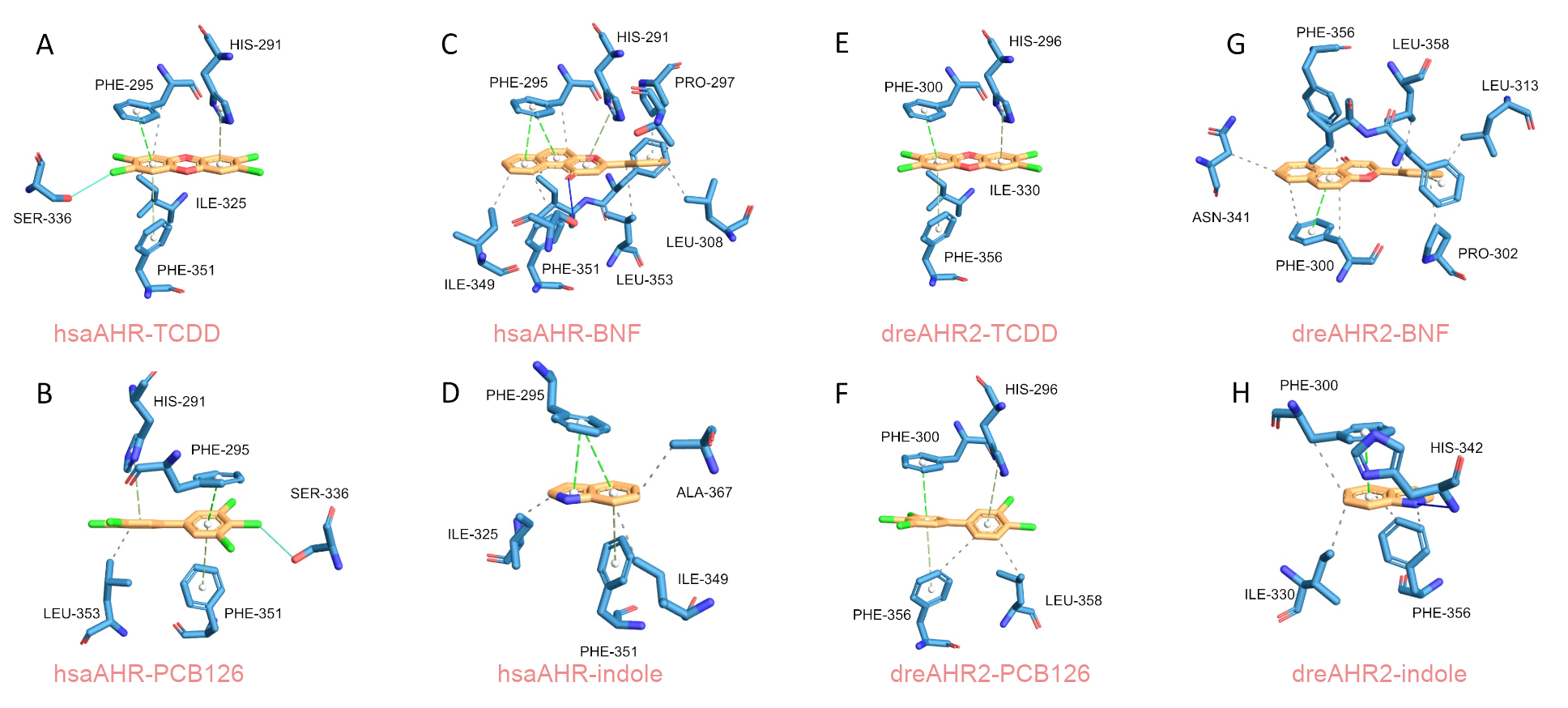

**Figure S4.** Interaction between hsaAHR/dreAHR2 LBD and ligands, including A (hsaAHR-TCDD), B (hsaAHR-PCB126), C (hsaAHR-BNF), D (hsaAHR-indole), E (dreAHR2-TCDD), F (dreAHR2-PCB126), G (dreAHR2-BNF), and H (dreAHR2-indole).

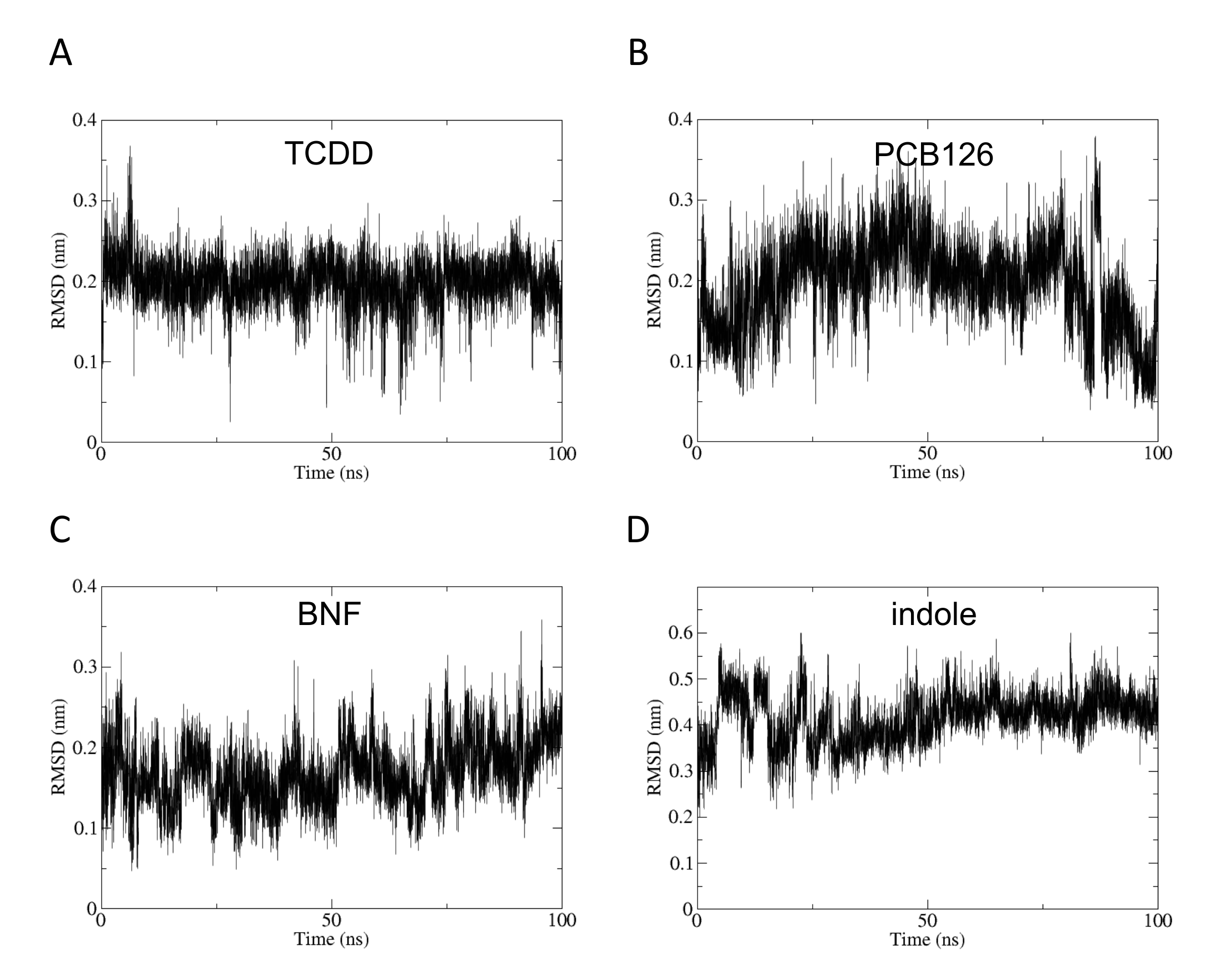

**Figure S5.** The RMSD of ligand-dreAHR2 interaction during 100ns simulation, including A (dreAHR2-TCDD), B (dreAHR2-PCB126), C (dreAHR2-BNF), and D (dreAHR2-indole).

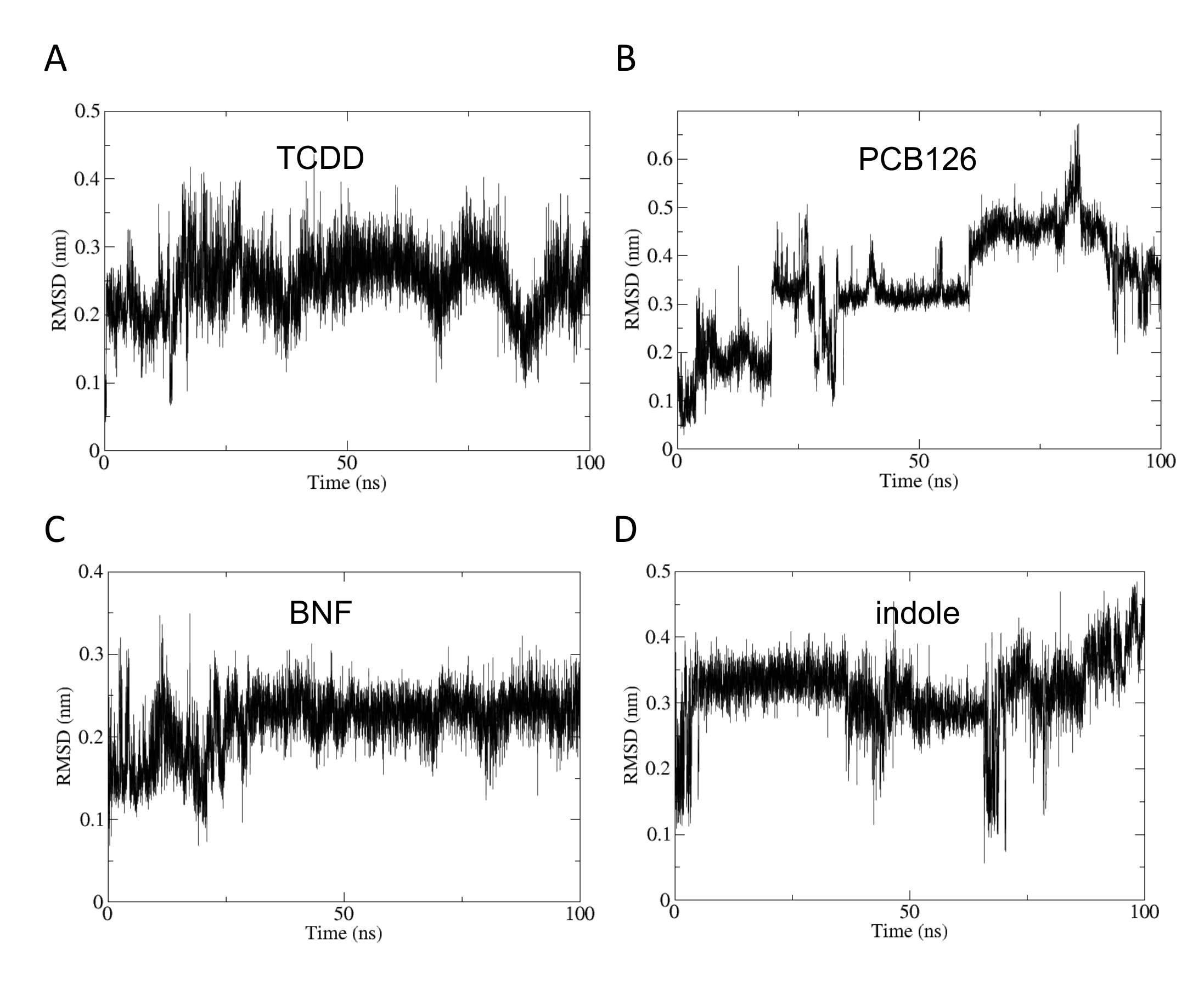

**Figure S6.** The RMSD of ligand-hsaAHR interaction during 100ns simulation, including A (hsaAHR-TCDD), B (hsaAHR-PCB126), C (hsaAHR-BNF), and D (hsaAHR-indole).
